# Biphasic mechanosensitivity of TCR mediated adhesion of T lymphocytes

**DOI:** 10.1101/232041

**Authors:** A. Wahl, C. Dinet, P. Dillard, P-H. Puech, L. Limozin, K. Sengupta

## Abstract

Force sensitivity of the T cell receptor (TCR) is now believed to be essential for immune recognition, but cellular mechanosensitivity of T cells is still poorly understood. Here we show that T cells adhering *via* the TCR-complex respond to environmental stiffness in an unusual biphasic fashion. As the stiffness increases, adhesion and spreading first increase, then decrease, attaining their maximal values on an optimally stiff surface, with stiffness comparable to certain antigen presenting cells. Remarkably, in presence of additional ligands for the integrin LFA-1, spreading increases monotonously with stiffness up to a saturation value. Using a mesoscopic semi-analytical model linking spreading to molecular characteristics of bonds, we identify force sensitivity of the off-rate and the effective bond stiffness as the crucial parameters that determine monotonic or biphasic mechanosensitive behavior.

---

Mechanosensitivity has emerged as a hallmark of many biological systems – from molecular to tissue level, and is implicated in health and disease (1–3). Soft and deformable substrates are used to study the impact of environmental stiffness on cells and to measure cell-generated forces. Pioneering single cell studies showed that for fibroblast cells grown on gels of various stiffness, locomotion and focal adhesions are regulated by substrate flexibility (4). Mechano-sensitivity has now been demonstrated for almost all cell types. Readouts include extent of spreading, cell stiffness, motility, differentiation and traction forces (5–7), and all typically increase monotonically with substrate stiffness, usually reaching a saturation value at a particular value of the stiffness. From a theoretical point of view, adhesion and mechanosensing can be treated either within cell-scale macroscopic models (3) that usually predict a monotonous increase in cell spreading and traction with stiffness, or using microscopic models that account for molecular mechanosensitivity (5), and may predict biphasic behavior in force or velocity (9). Experimental reports of such biphasic behavior is very rare for cells adhering to a surface, examples include speed of migrating neutrophils (10), actin retrograde flow (5) and traction forces in beating muscle cells (11) or talin-silenced fibroblasts (12), the adhesion area still being monotonous whenever reported.

A vast majority of experimental studies on mechanotransduction are conducted on focal adhesion forming cells. Leucocytes in general and lymphocytes in particular have been much less studied, despite evidence that T cells response is sensitive to substrate stiffness (13, 14), but contradictory trends were been reported (15, 16), which probably hinted at a non-monotonicity. The ability of T cells to recognize pathogens and pathological cells is now known to depend on their mechanosensitivity (13,14), which implicates the CD3 domain of the T cell receptor (TCR) complex (17). Here we explore mechanosensing in T cells *via* the TCR-CD3 complex, in presence or absence of the ligand ICAM-1 for the T cell integrin LFA-1. We follow the spreading of Jurkat T cells, which on this cell type is a marker of activation (18), on functionalized surfaces of polydimethylsiloxane-based (PDMS silicone) elastomers with stiffness (quantified in terms of Young’s modulus) ranging from 500 Pa to several MPa. To cover such a large range while retaining similar surface chemistry, different silicone types with adapted base/cross-linker ratio were prepared and characterized in terms of their Young’s modulus from force curves obtained using atomic force microscopy nano indentations. The PDMS surface was then functionalised with either anti-CD3 alone with estimated molecular coverage of about ~ 400 moleculesμm^2^, or, additionally with ICAM-1 at similar coverage (Table S1).

In a first set of experiments, T cells were allowed to interact with anti-CD3 functionalized surfaces (Fig. 1A) for twenty minutes, allowing them enough time to spread but not retract. They were then fixed and stained for actin, TCR or the zeta-chain-associated protein kinase 70 (ZAP-70–one of the first molecules to be recruited to the TCR complex upon TCR engagement). Cells were imaged in bright field, reflection interference contrast microscopy (RICM), total internal reflection fluorescence (TIRF) and confocal microscopy (Fig. 1B and fig. S1). As can be immediately seen, cells spread more on 20 kPa than on 2440 kPa elastomers but the actin is peripherally distributed in both cases. RICM images were analyzed to quantify the cell spreading area (19–21) and the actin TIRF images to quantify the extent of actin peripherality. Fig 1C and fig. S2 summarize the final cell area measured for different elastomer stiffness. Note that for the same stiffness, changing the silicone type does not affect cell area. Cells on the softest elastomer (QGel 920 1:0.95 at 0.5 kPa) spread moderately to an area of about 200 μm^2^ The area increases as the stiffness is increased to 1 kPa (CY52 1:1) and then to 4 and 5 kPa (QGel 920 1:1.1 and Sylgard 184 1:58, called soft), reaching a maximum of about 300 μm^2^. Thereafter the cell spread area decreases with increasing stiffness, falling back to roughly 200 μm^2^ for 2 MPa (Sylgard 184 1:10, called hard) and to less than 150 μm^2^ at 7 MPa (Sylgard 184 1:10, rigidified by plasma-treatment). On equivalently functionalized glass, provided that non-specific interactions are fully blocked, here using ligands immobilized on supported lipid bilayers (21), the cells spread to a mere 120 μm^2^ (without full blocking, area on anti-CD3 coated glass is high at ~300 μm^2^). This remarkable spreading behavior is reproduced in human primary CD4+ T cells, which spread more on soft than on hard PDMS substrates (fig. S3).

**Fig. 1.**
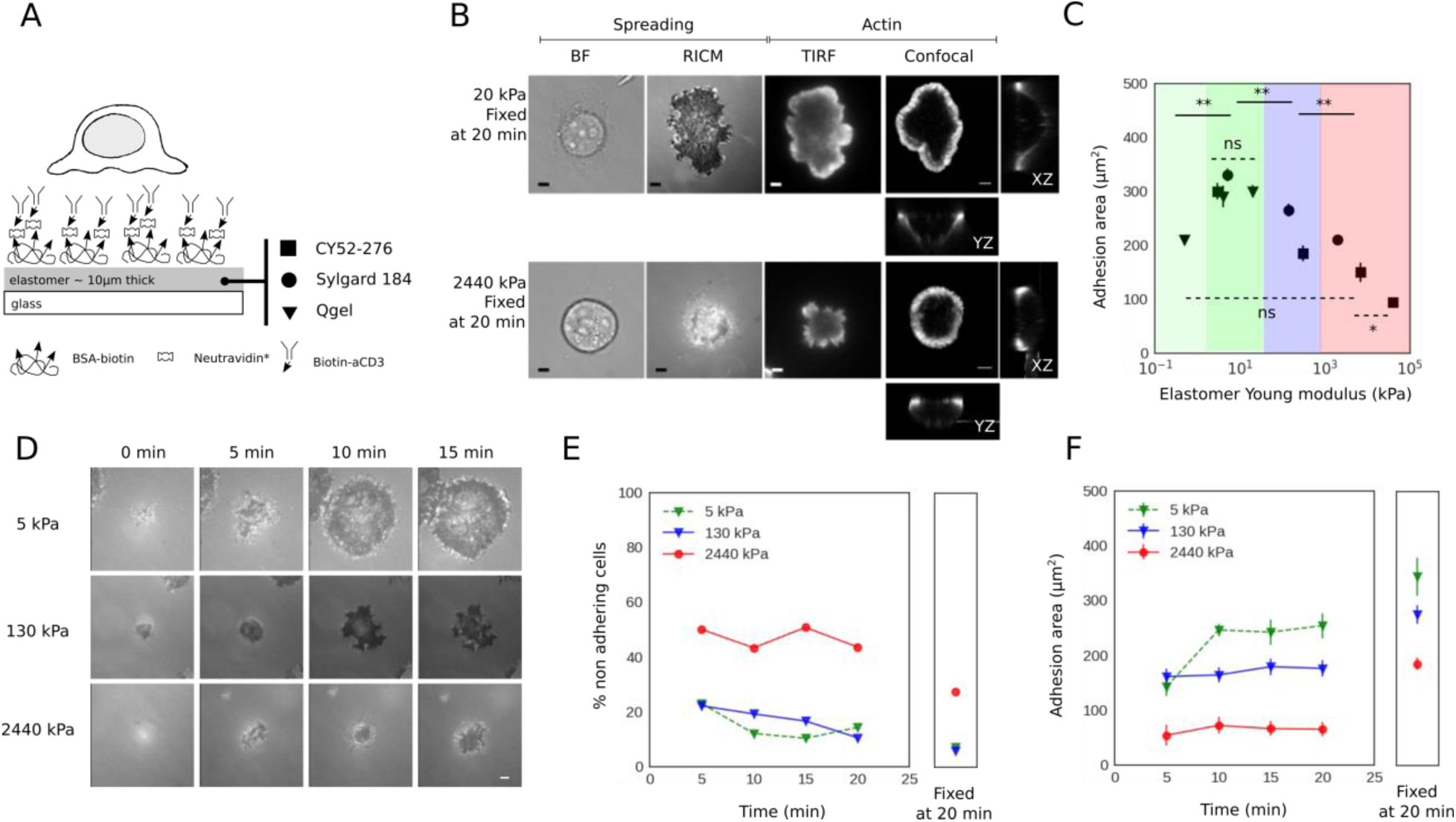
T cell spreading on elastomers functionalized with anti-CD3. (A) Schematic representation of the experiment. T cells interact with elastomers with stiffness ranging from 500 Pa to 40 MPa, and functionalized with anti-CD3. (B) T cells were allowed to spread for 20 minutes on either soft (20 kPa) or hard (2440 kPa) substrates, were fixed and imaged in BF, RICM, TIRF (actin) and confocal (actin) modes. (C) Cell spread area as quantified from RICM images. Each point is average of at least 50 cells and 3 samples (see fig. S2 for statistical tests). The range of very soft (light green), soft (green), intermediate (blue) and hard (red) are color-coded. (D) Time course of cell adhesion imaged in RICM. (E) Percentage of non-adherent cells as time progresses, and after fixation. (F) Quantification of adhesion area of the adhered fraction. At least 100 cells for each time point and each stiffness. Note that weakly adherent cells are washed away during fixation, thus driving the average area towards higher values. Throughout, error-bars are SEM. n.s. indicates no significant difference. Scale bar 4 μm.

T cells tend to spread isotropically on functionalized glass resulting in a roughly circular shape, whereas on the elastomers studied here, they may exhibit irregular star-like shapes (fig. S4). The shape of a cell is mainly determined by its actin cytoskeleton. The distribution of actin at the adhesive interface of a fully spread T cell tends to be peripheral, but the peripherality becomes less pronounced in weakly spread cells (21, 22). Here, a peripheral ring-like distribution is seen for all stiffness values (Fig. 1B, fig. S1, quantified in fig. S5), even when the cell area is relatively small. The recruitment of TCR and ZAP-70 kinase imaged in TIRF do not show appreciable variations with stiffness (fig S1), hinting at a mechanistic rather than signalling based mechanosensing mechanism.

We next explored the dynamics of T cells on soft (5 kPa), intermediate (130 kPa) and hard (2.5 MPa) substrates. Fig. 1D shows an example of single cell time-lapse RICM demonstrating that the cells on hard substrates lag behind in spreading already in the time window 0 to 5 minutes, a period shown earlier to be critical for antigen recognition (18). Fig. 1E and F quantify this effect at the scale of the population. It is seen that on hard substrates there is a population of nonadherent cells that never adhere (Fig. 1E). Furthermore, the cells that do adhere, spread much less on the hard substrate (Fig. 1F). The time evolution of the actin organization is similar in the two cases (fig. S6).

In the next set of experiments, the role of LFA-1 was explored by dual functionalization of the substrates with anti-CD3 and ICAM-1. As negative control, and consistent with past reports of cells on glass supports with only ICAM-1 on surface (21), there is no adhesion on soft (5 kPa) or hard (2MPa) PDMS (fig. S7). Interestingly, this is also the case when cells are simultaneously stimulated with anti-CD3 in solution. For the case of co-functionalization with both anti-CD3 and ICAM-1 (Fig. 2A), the cells spread the least on the softest substrates used and they spread more as the substrate stiffness is increased, already reaching a saturation value of about 400 μm^2^ in ~kPa range (earlier designated as soft, Fig. 2B,C). Thus the maximal area reached here is somewhat larger than that with anti-CD3 alone at 300 μm^2^. Analysis of time-lapse RICM data shows that the spreading dynamics does not differ between the 5 kPa and 2.5 MPa cases (Fig. 2D). In terms of actin organization, no clear difference is detected between soft and hard on one hand and with or without ICAM-1 on the other hand; the actin is mainly peripheral in all cases (fig. S5).

**Fig 2.**
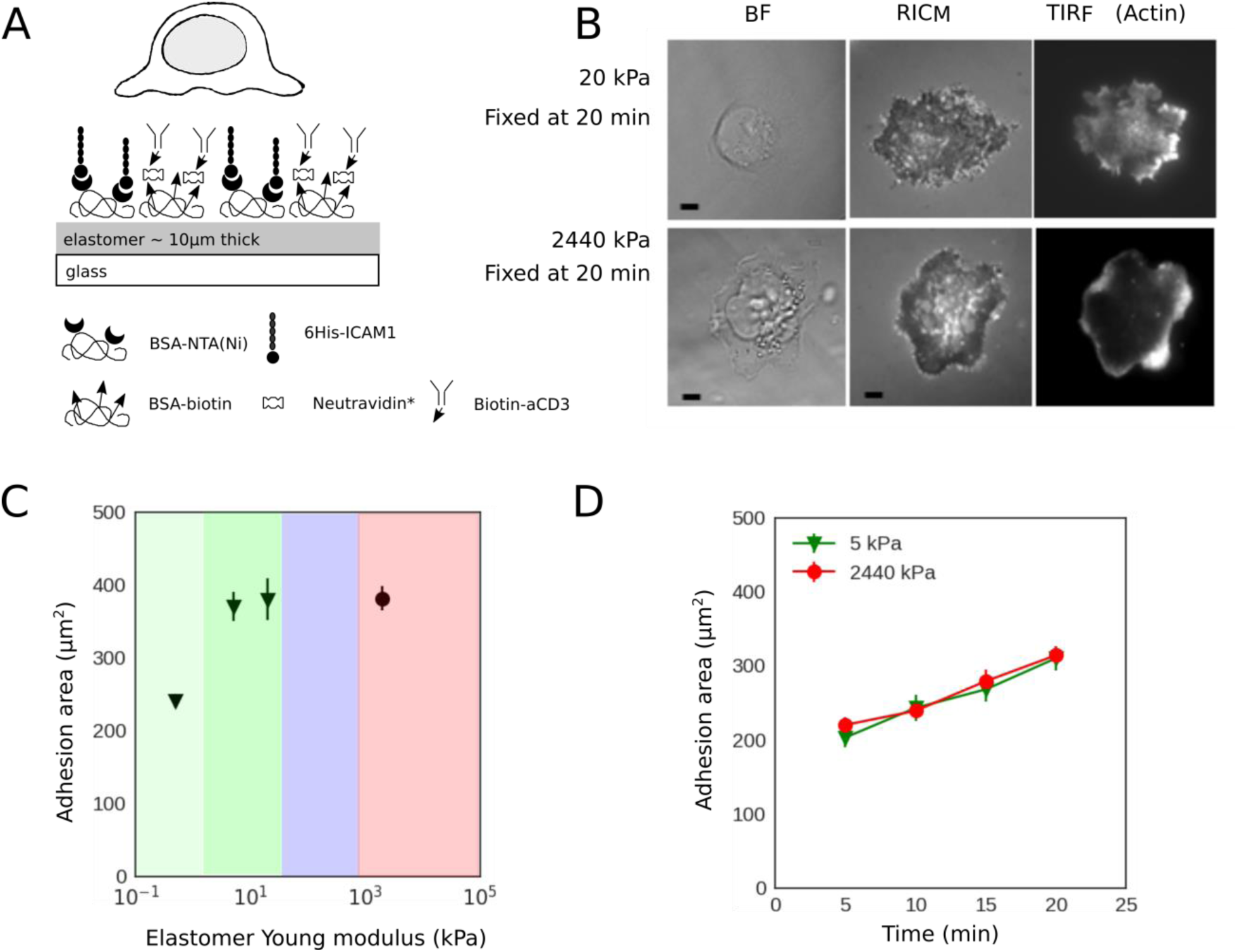
T cell spreading on elastomers co-functionalized with anti-CD3 and ICAM-1. A. Schematic representation of the experiment. T cells interact with elastomers with stiffness ranging from 500 Pa to 2.5 MPa, which are dually functionalized with ICAM-1 and anti-CD3. B. T cell spreading on such substrates imaged in BF, RICM and actin in TIRF after 20 min of spreading and fixation. C. Corresponding cell spread area, averaged over at least 50 cells and at least 3 samples per data point. D. Adhesion area as a function of time. Mean values are reported (at least 100 cells for each time point and each stiffness, average area is slightly over-estimated due to 30 to 40 nonadherent cells ignored for the analysis). Error bars are SEM. Scale bar 4 μm.

The spreading behavior is strikingly reproduced in human CD4+ naive T cells isolated from peripheral blood, which fail to spread on anti-CD functionalized hard PDMS without ICAM-1 but do spread on hard PDMS dually functionalized with ICAM-1 and anti-CD3, as well as on soft PDMS with or without ICAM-1 (fig. S3).

We next prepared soft, intermediate and hard substrates, dually functionalized with anti-CD28 and anti-CD3 in equi-molar ratio, to assess the impact of engaging the co-stimulatory molecule CD-28. Interestingly, the presence of anti-CD28 has no impact on the cell area, which remains unchanged with respect to the case without anti-CD28 (fig. S8). This is consistent with earlier work suggesting T cell mechanosensing is actuated through CD3 rather than CD28 (17). Since acto-myosin generated forces are thought to be at the heart of mechanosensing, we tested the effect of inhibiting myosin IIA activity through the drug blebbistatin. It was shown earlier that in case of cells that spread weakly on mobile ligands, myosin treatment rescues spreading (21). However, surprisingly, no effect of myosin IIA inhibition was observed here (fig. S9).

The lack of a role for myosin, a major player in traditional clutch based models (8), led us to identify actin polymerization generated forces as the basis for theoretical description of mechanosensitive spreading. In the past, mesoscopic models have linked actin polymerization driven cellular scale spreading (21) or traction force generation by a filopodium pulled by optical tweezers (23), to force dependence of bond kinetics. Here we couple the two approaches to model the edge of the cell where lamellipodia-like protrusions get pushed forward by actin polymerization just behind the membrane (Fig 3A). Within a quasi-one-dimensional model of such a cell edge, the actin is modelled as a strip which polymerizes at a given velocity, and pushes the frontier of the cell forward, at the same time generating a retrograde flow of the actin away from the edge with a velocity *v_p_* (Fig 3B). The final cell spreading area (*A*) is then set by a balance of the actin generated force (*F*, expressed as force density) and a tensile force that limits spreading, such that *F* is proportional to *A* (see Supplementary Materials).

**Fig 3.**
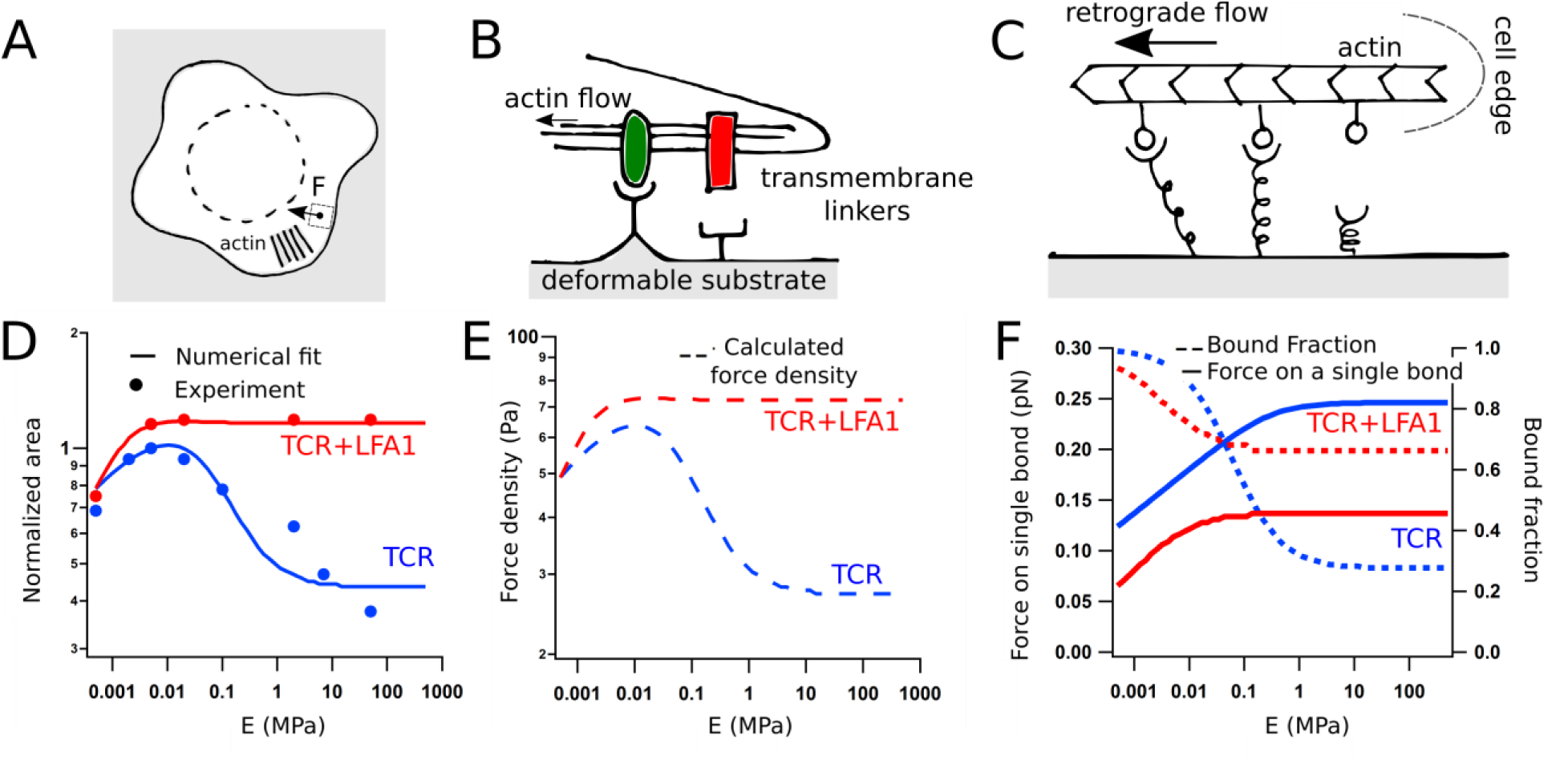
The model and fit to data. A. Schematic representation of the model of T cell spreading at the cell scale. B. Mesoscopic view where cells interact with elastomers through bonds of one or two types, that link the actin to the substrate. C. The entire actin-receptor-ligand-substrate connection is represented as a single effective linker. D. Fit of model to normalized area data. E. Force density in Pascal. F. Force on a single bond and the fraction of receptors that are bound.

The ligands are tightly bound to the elastic substrate and form bonds with cellular transmembrane receptors, whose intracellular moieties interact with actin *via* adapter proteins that are mostly known for integrins but are still a matter of debate for TCR. For simplicity, the entire complex is modelled here as a single spring-like bond (henceforth called a linker, Fig. 3C). *F* is transmitted to these bonds such that the force *f* on a single bond is *F*/(*nN*) where *n* is the bound fraction, and *N* the ligand density. The bond kinetics is defined by a constant on-rate (*k_on_*) and a force dependent off-rate given by 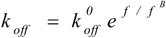 where 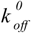 is the off-rate at zero force and *f ^B^* is the characteristic Bell force at which a bond becomes force sensitive. The linker has an intrinsic bond elasticity *k_b_* which is renormalized by the substrate elasticity (*E*) in such a way that it increases with *E*, reaching a saturation when the value of *E* exceeds *kb*/*a, a* being a molecular length scale (23). Note that for anti-CD3 alone (henceforth called TCR case), the linker characteristics may be measurable from single-molecule experiments. In presence of ICAM-1 (henceforth called TCR+LFA-1) however, a single effective linker represents both TCR and LFA-1 mediated bonds – the attributed molecular parameters are effective quantities that embody the real parameters corresponding to each linker type as well as possible cooperative effects.

The model self-consistently determines *F* and *n*, for given *N* and *v_p_* values which were measured independently. A fit of the model to area data is able to capture the experimentally observed stiffness dependence (Fig. 3D). For a given *v_p_*, both *F* and *f* initially increase with *E via* the renormalization of *k_b_* (Fig. 3E,F). However, when *f* becomes appreciably larger than *f ^B^*, bonds start breaking faster than they form and *n* drops, leading to a drop in *F* at *E*>*E*\*, even though *f* continues to increase with *E*. In the TCR case, the bonds are predicted to be stiff and very sensitive to force (high *k_b_* and low *f ^B^*), ensuring that *k_off_* overtakes *k_on_* before *E* reaches *kb*/*a*. In the case of TCR+LFA1 however, *k_b_* saturates before *E*\* can be attained. The existence of *E*\* is an indicator of whether or not the force, and therefore the area, exhibits biphasic behavior. Fig. S10 show plots of *E*\* as a function of the parameters *k_b_*, *f ^B^*, and 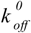 (keeping *k_on_* constant). It is again seen that independent of 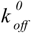, high *k_b_* and low *f ^B^* are essential for biphasic behavior. From the fits, the value of *f ^B^* is expected to be about 0.2 pN for TCR and about the double for TCR+LFA1.

Previous single-molecule probing of TCRs on T cells (24) yielded different force response compared to cell-free experiments with purified TCRs (25). Furthermore, the characteristic force *f ^B^* for a typical ligand-receptor bond is expected to be in the pN range. This leads us to speculate that the highly force-sensitive bond described above for the TCR case is in fact the previously mooted CD3 to actin link (21, 26), rather than the ligand-receptor (anti-CD3/TCR) bond. On integrin engagement, this putative link is probably reinforced and the effective link is less force sensitive. Interestingly, measurements in the range of elasticity spanning up to few kPa do report higher traction forces and activation on stiffer substrates (13, 15, 27). This range, roughly coinciding with the first rising phase of our biphasic curve, is also the physiological range of elasticity for professional antigen presenting cells, which may be modified under pathological conditions (28). Whether T cells exploit the potentially biphasic response of the TCR-CD3 complex for their physiological function remains to be explored, but the putative CD3-actin link proposed here as highly force sensitive should inform future studies on T cell mechanoresponse.

## Acknowledgments

We thank Anne Charrier for help and guidance for elastomer stiffness measurements and critical reading of the manuscript, M. Biarnes-Pelicot for cell culture and isolation, CINaM-PLANET for profilometer measurements, the PCC culture facility and L. Borge for cells, Anais Sadoum for complementary experiments, Emmanuelle Bouveret for generous gift of YFP-his and Pierre Sens for fruitful discussions. We also thank Pierre Bongrand for stimulating discussions and critical reading of the manuscript. This work was partially funded by European Research Council via grant no. 307104 FP/2007-2013/ERC-Stg SYNINTER. Additional data and text can be found in the Supplementary Materials.

